# Alcohol induces p53-mediated apoptosis in neural crest by stimulating an AMPK-mediated suppression of TORC1, S6K, and ribosomal biogenesis

**DOI:** 10.1101/2024.07.02.601754

**Authors:** Yanping Huang, George R. Flentke, Susan M. Smith

**Affiliations:** UNC Nutrition Research Institute, University of North Carolina at Chapel Hill, Kannapolis NC USA; Dept. Nutrition, University of North Carolina at Chapel Hill, Kannapolis NC USA

**Keywords:** AMPK, apoptosis, craniofacial development, fetal alcohol spectrum disorders, mTOR, neural crest, nucleolar stress, O9-1 cells, p53, RPS6K, ribosome biogenesis

## Abstract

Prenatal alcohol exposure is a leading cause of permanent neurodevelopmental disability and can feature distinctive craniofacial deficits that partly originate from the apoptotic deletion of craniofacial progenitors, a stem cell lineage called the neural crest (NC). We recently demonstrated that alcohol causes nucleolar stress in NC through its suppression of ribosome biogenesis (RBG) and this suppression is causative in their p53/MDM2-mediated apoptosis. Here, we show that this nucleolar stress originates from alcohol’s activation of AMPK, which suppresses TORC1 and the p70/S6K-mediated stimulation of RBG. Alcohol-exposed cells of the pluripotent, primary cranial NC line O9-1 were evaluated with respect to their S6K, TORC1, and AMPK activity. The functional impact of these signals with respect to RBG, p53, and apoptosis were assessed using gain-of-function constructs and small molecule mediators. Alcohol rapidly (<2hr) increased pAMPK, pTSC2, and pRaptor, and reduced both total and pS6K in NC cells. These changes persisted for at least 12hr to 18hr following alcohol exposure. Attenuation of these signals via gain- or loss-of-function approaches that targeted AMPK, S6K, or TORC1 prevented alcohol’s suppression of rRNA synthesis and the induction of p53-stimulated apoptosis. We conclude that alcohol induces ribosome dysbiogenesis and activates their p53/MDM2-mediated apoptosis via its activation of pAMPK, which in turn activates TSC2 and Raptor to suppress the TORC1/S6K-mediated promotion of ribosome biogenesis. This represents a novel mechanism underlying alcohol’s neurotoxicity and is consistent with findings that TORC1/S6K networks are critical for cranial NC survival.

## 1. Introduction

Prenatal alcohol exposure (PAE) is a leading cause of permanent neurodevelopmental disability and features deficits in cognition and executive function [1]. Diagnosis is often initiated by a distinctive craniofacial appearance [2] that has complex origins reflecting both changes in the underlying brain growth [3,4] and direct effects upon the cranial neural crest [5,6], a pluripotent stem cell linage that forms the facial bone and cartilage, and neuronal elements including certain cranial nerves, Schwann cells, and the sympathetic and parasympathetic system [7]. Neural crest induction begins at neurulation and, shortly thereafter, they emigrate from the closing dorsal neural tube and into peripheral tissues where they differentiate into the aforesaid structures. Pharmacologically relevant alcohol exposure (10-80mM) disrupts multiple events in cranial neural crest development including their induction, migration, proliferation, and survival [4–6]. We and many others have shown that alcohol causes the apoptotic deletion of neural crest progenitors at a specific phase of their development, at the onset of migration [8–10]. Alcohol interacts with multiple pathways to modulate facial outcome and/or neural crest death including sonic hedgehog, retinoid, oxidative stress, Wnt, and p53, among others [11–17]. However, the regulatory mechanisms governing neural crest sensitivity to alcohol remain elusive. Novel insight into this apoptotic mechanism emerged from our whole exome sequencing of neural crest progenitors, wherein a brief alcohol exposure (2hr, 52mM) rapidly (<6hr) repressed the expression (*p_adj_*=10E-47) of 106 nuclear and mitochondrial ribosomal proteins (RP) by 30%-50% [18]. A similar repression of RPs is observed in alcohol-exposed zebrafish [19], in the headfolds of mouse [20,21] and chick strains [22] having differential alcohol vulnerability, in primary mouse neural stem cells [23] and in the cranial neural crest primary cell line O9-1 [24]. Thus, suppression of RP synthesis is a consistent feature of alcohol-exposed neural crest and neuroprogenitors across taxons.

Why would a suppression of RPs lead to the apoptosis of cranial neural crest populations? For rapidly proliferating populations such as neural crest, each cell division requires the duplication not only of DNA, but the ribosomes that accomplish protein synthesis. Indeed, the production of rRNA plus the ∼200 proteins and short RNAs necessary for rRNA processing and assembly into ribosomes is estimated to occupy 70-80% of the cellular energy budget [25]. This process is so abundant that it is seen visually in the nucleosomes, which are the multiple operon sites of this ribosome biogenesis (RBG) [26]. Unsurprisingly, cells have coopted RBG to monitor their internal stress, and RBG is tightly coupled to cellular anabolism and the p53 checkpoint pathway [27,28]. Specifically, under normal conditions RBG is stimulated by the anabolic effector TORC1 via S6K [28]. When energy is limiting, pAMPK phosphorylates the TORC1 components TSC1/2 and Raptor to suppress TORC1 and RBG, thus integrating RBG with cell metabolism. Under stressors that reduce the resources needed for rRNA synthesis, RBG ceases and the nucleoli visually disappear [29,30]. This process, known as “nucleolar stress”, is linked to p53 via the nuclear E3 ubiquitinase MDM2, which under normal conditions targets and destabilizes p53 [31–33]. Under nucleolar stress, reduced rRNA synthesis enables a RPS5-RPL11-5SrRNA complex to bind and inhibit MDM2, thus allowing p53 to become transcriptionally active [31–33]. The importance of RBG for neural crest is evidenced in the genetic disorders known as ribosomopathies, in which loss-of-function in AMPK-TORC1-S6K signaling, or in components of RBG such as rRNA, RPs, or the ribosome assembly machinery, cause p53-mediated neural crest losses and facial deficits that share similarities with those of PAE (i.e. Treacher-Collins syndrome, Diamond-Blackfan anemia) [34–36].

We recently reported that alcohol exposure impairs RBG to induce nucleolar stress in early cranial neural crest progenitors. Alcohol exposure causes the rapid cessation of rRNA synthesis and dissolution of nucleolar structures in response to pharmacologically relevant alcohol exposures (20-80mM for 2hr) [24]. This is followed by the stabilization of nuclear p53 and their subsequent p53-mediated apoptosis [17,24]. Haploinsufficiency of RPL5, RPL11, or RPS3A using a morpholino approach synergizes with alcohol to cause craniofacial deficits in a zebrafish model of PAE [18,24], and transfection with MDM2 or loss-of-function p53 blocks these effects of alcohol [24]. However, the mechanism responsible for alcohol’s suppression of RBG is unknown. Here we test the hypothesis that alcohol represses RBG through its suppression of S6K via TORC1, and that this activation of AMPK contributes to both the loss of TORC1/S6K signaling and their p53-mediated apoptosis in response to alcohol exposure.

## 2. Materials and Methods

### 2.1. Cell Culture

All studies use the established primary cranial NC line O9-1 (#SCC049, Millipore; Burlington, MA) [37,38]. This non-transformed stem cell line was originally isolated from mass culture of primary cranial NCs isolated at embryonic day (E)8.5 from C57BL/6J mouse embryos that expressed Wnt1-Cre:R26R-GFP [37]. It expresses key NC markers (e.g., Twist1, Snail1, Nestin, CD44), can differentiate into osteoblasts, chondrocytes, smooth muscle cells, tooth pulp, and glial cells, but not neuronal cells, and develop normally when added back to embryos [37–40]. We find it has alcohol responses identical to those of primary avian or mouse NCs [24]. Cells were maintained on Matrigel-coated plates in their pluripotent state at 50-60% confluence in DMEM (Gibco, Grand Island NY) supplemented with 6% ES cell media (ES-101B, Sigma, Louis MO), 15% fetal calf serum (Millipore), 0.1mM nonessential amino acids (Gibco), 1000 U/ml leukemia inhibitory factor (LIF, Gibco), 25ng/ml basic fibroblast growth factor (Invitrogen, Waltham, MA), 0.3µM β-mercaptoethanol (MP Biomedical, Solon, OH), and 100 U/ml penicillin and streptomycin (Gibco). Experiments were performed in the identical media but containing 1% FCS.

### 2.2. Alcohol Exposure

Most studies used a 2 x 2 experimental design (Control (CON) vs. Alcohol (ALC); ± Intervention) in which all groups originated from the same starting culture and were incubated and harvested simultaneously. Cells were cultured overnight, and then the culture media was removed and replaced with either normal media or media containing the experimental intervention. After one hour, the media was removed, the cells briefly washed, and then incubated with media-only (CON) or media containing 80mM ethanol (ALC, 100%, USP grade; Koptec; King of Prussia, PA). We selected this concentration because we previously found this is the EC50 for the alcohol-mediated dissolution of nucleolar structures in O9-1 cells [24]. In these culture conditions, we find that the alcohol concentrations drop to less than 10mM by 2hr post-addition. Thus, these studies model acute binge exposure.

### 2.3. Nucleolar Immunostaining

Control and alcohol-exposed O9-1 cells were generated as above and fixed in 4% paraformaldehyde at 4hr thereafter. Immunostaining for the nucleolar protein upstream binding factor (UBF; Abcam #61205, 1:250) was performed exactly as described in [24]. Images were captured under uniform exposure.

### 2.4. Whole Transcriptome Sequencing

Control and alcohol-exposed O9-1 cells were generated as above, and 6hr thereafter were harvested using TrypLE Express (Gibco) as per manufacturer’s directions and flash-frozen. Cell pellets containing ∼10^6^ cells were submitted to the UNC High Throughput Sequencing Facility for cDNA library preparation and whole transcriptome sequencing as described previously [23]. Sequence quality, read alignments, and gene assignments were performed as described [23]. Differential expression was analyzed using DESeq2 and adjusted for false discovery using the Bonferroni method [23]. Expression-level changes were considered significant at an adjusted p-value ≤0.05.

### 2.5. Quantitative PCR

RNA was isolated using Trizol reagent (Invitrogen) from CON and ALC cultures at experimentally determined times following exposure to 80mM alcohol. Total RNA (1µg) was reverse transcribed with 500µg/ml random primer (#C1181), 5X ImProm II reaction buffer (#M289A), ImProm II (#M314A), 25 mM MgCl_2_ (#A351H), 10 mM dNTP (#U1511), 20U RNAsin (#N2511; all from Promega, Madison, WI) and RNAse free water. Quantitative PCR (qPCR) were performed using SYBR Select Master Mix (ABI, #4472913) and the Real-Time PCR system (Bio-Rad CFX96; Hercules, CA). *De novo* rRNA synthesis was quantified by qPCR for the Internal Transcribed Spacer ITS1 (138bp) located between the 18S and 5.8S rRNA of newly transcribed 47S rRNA [23,24]. The primers (IDT, Coralville, IA) for ITS-1 were forward: 5′-CCGGCTTGCCCGATTT-3′, reverse: 5′-GCCAGCAGGAACGAAACG-3′ and for β2 microglobulin were forward: 5′-TTCACCCCCACTGAGACTGAT-3′; reverse: 5′-GTCTTGGGCTCGGCCATA-3′. Relative abundance was calculated using the 2^−ΔΔCT^ method and was normalized to β2 microglobulin. qPCR methodology adhered to the MIQE standards [41].

### 2.6. Western Immunoblot Analysis

Total proteins were isolated from cells at the indicated times using NP-40 lysis buffer (50 mM Tris-HCl, pH 7.5, 150 mM NaCl, 0.5% NP-40, 50 mM NaF, 1 mM NaVO_3_, 1 mM dithiothreitol, 1 mM phenylmethylsulfonyl fluoride; all from Sigma). Total protein was separated on 10% SDS-PAGE reducing gels for proteins ≤150kDa, or 4%-20% detergent gels for those ≥150kDa (BioRad). The resolved proteins were transferred to a PVDF membrane using the Trans-Blot Turbo Transfer System (Bio-Rad). Primary antibodies directed against p-AMPK(Thr172) (#2535), AMPK (#2793), PKCα (#2056), pRaptor(Ser792) (#2083), Raptor (#2280), pRictor(Thr1135) (#3806), Rictor (#2140), RPS6 (#2217), pRPS6(Ser235) (#2211), pS6K1(Thr389) (#9234), S6K1 (#9202), pTSC2(Ser1387) (#5584), TSC2 (#4308), p4E-BP1(Thr70) (#9455), and 4E-BP1(#9452) were all used at 1:1000 and were from Cell Signaling (Danvers MA). Antibodies directed against p53 (#ab26, 1:500), Deptor (#191841, 1:200) and pPKCα(Ser657) (#ab180848, 1:500) were from Abcam (Cambridge, England, UK). The secondary antibodies were goat anti-rabbit (#4010-05, 1:5000) and anti-mouse immunoglobulin (#1030-05, 1:5000, both from Southern Biotech, Birmingham AL) coupled to horseradish peroxidase. Signal was imaged by chemiluminescence using the Radiance Q detection system (Azure Biosystems, Dublin, CA), and data were normalized to total protein per lane using Revert 700 Total Protein Stain (Licor, Lincoln NE).

### 2.7. Cell Transfection

Cells were transiently transfected at 80% confluence with plasmid pcDNA3 S6K2 E388 D3E (2.5 µg/µl; #17731; Addgene; Watertown, MA; 42) or the control empty plasmid pCS2 (2.5 ug/ul) and Lipo-3000 (#L3000001; Invitrogen) according to the manufacturer’s instructions. This construct contains four gain-of-function mutations (T388E, S410D, S417D, S423D) that confer rapamycin-resistance and constitutive activity (caS6K) [42]. Cells were incubated with plasmid and Lipo-3000 for 36hr, washed, and then reincubated for 6hr prior to starting the alcohol exposure.

### 2.8. Apoptosis Analysis

Cells were exposed to 80 mM alcohol as per above and fixed 16hr thereafter in 4% paraformaldehyde, permeabilized in 0.2% TritonX-100, and apoptosis was detected using TUNEL assay (In Situ Cell Death Detection kit, TMR Red; Roche, Indianapolis, IN). Nuclei were visualized using DAPI. Images were taken under uniform exposure, capturing three images per technical replicate.

### 2.9. Small Molecule Interventions

These used the AMPK inhibitor dorsomorphin (5µM; IC50 = 1µM), the AMPK activator A769662 (100µM; EC50=6 µM), the TORC1 inhibitor rapamycin (1 nM; IC50=00.1nM; all from Selleck Chemicals, Houston TX), or the TORC1 activator L-leucine (250 µM, Sigma; media concentration 67 µM). Experimental concentrations were based on review of the literature and were within an order-of-magnitude of the IC50 or EC50, taking into account bioavailability. All compounds were dissolved in water except for rapamycin, which was first dissolved in DMSO; these stocks were then added to bulk media at 1:10,000). For all, cells were incubated 1h in media containing the compound or media-only, washed, and then treated with fresh media or media containing 80mM alcohol, and collected at experimentally determined times thereafter.

### 2.10. Statistical Analysis

All data are the mean ± SD of three independent experiments, each experiment having three replicates per group unless indicated otherwise. Data were tested for normality and then analyzed using the appropriate test, Student’s t-test for comparisons of two groups and one-way or two-way ANOVA for multiple comparisons. *P* < 0.05 was considered significant.

## 3. Results

### 3.1. Alcohol suppresses RBG and induces nucleolar stress and p53-mediated apoptosis in neural crest

As we showed previously [24], a time course reveals a significant interaction between alcohol exposure and time (F(7,28)=4.948, p=0.001) such that exposure to 80mM alcohol suppressed *de novo* rRNA synthesis by 1hr following exposure (CON 100 ± 11.7%, ALC 63.5 ± 4.8%; *p*=0.007; **Figure 1A**), and this reduction persisted for at least 4hr thereafter (*p*=0.024). Also as shown previously [24], this was followed by 4hr following the alcohol exposure by the dissolution of nucleolar structures that was visualized, for example, using antibodies directed against the RNA polymerase I transcription factor UBF (**Figure 1B**) [24,29]. The proapoptotic protein p53, which responds to nucleolar stress via MDM2 [31–33], was stabilized in these cells within 2hr of exposure, when its content increased to 170 ± 11% of CON values (*p*<0.001). We showed previously [24] that this largely represents nuclear protein and p53 remained elevated for at least 18hr following the alcohol exposure (191 ± 14% of CON; *p*<0.001; **Figure 1C**). By 16hr, the alcohol-exposed neural crest underwent apoptosis (CON 0.97 ± 0.23%, ALC 10.94 ± 1.24%; *p*<0.001; **Figure 1D**), and we have shown elsewhere that p53 mediates this death [24].

**Figure 1.**
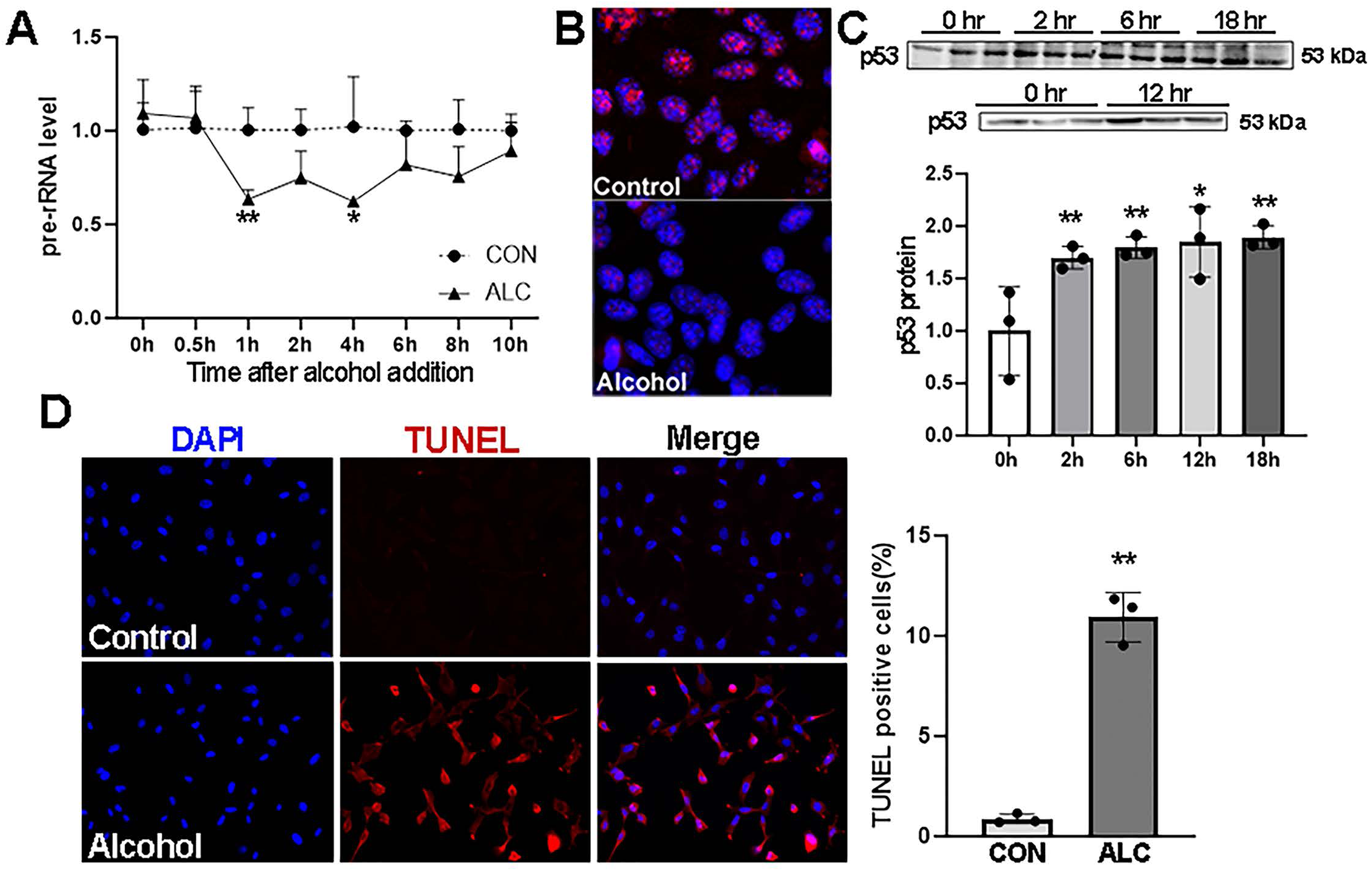
Alcohol rapidly represses ribosome biogenesis and causes nucleolar stress and p53-mediated apoptosis in primary neural crest cells. **(A)** ITS-1, a product of de novo rRNA synthesis, is reduced by 1hr following exposure to alcohol. **(B)** Nucleoli in neural crest are reduced and become diffused within 4hr of alcohol exposure, as visualized using immunostain for the nucleolar protein UBF. **(C)** p53 is stabilized in ALC neural crest within 2hr of exposure and this persists for at least 18hr. **(D)** Alcohol causes apoptosis in neural crest at 16hr post-exposure. Red signal is TUNEL, blue is DAPI stain to visualize nuclei. All studies use 80mM alcohol. Values are mean ± SD of N=3 independent experiments. * p<0.05, **p<0.01, ***p<0.001 compared with CON by either two-tailed T-test (D) or one-way (C) or two-way ANOVA (A). CON, control; ALC, alcohol-exposed.

### 3.2. Alcohol exposure reduces S6K, and S6K gain-of-function normalizes rRNA synthesis and apoptosis in neural crest

To gain insight into the mechanism by which alcohol induced nucleolar stress in neural crest, we evaluated its impact upon a primary effector of RBG, S6K [28]. Alcohol exposure reduced the abundance of S6K in a complex manner. With respect to total S6K protein, its content was 69 ± 6% (*p*<0.001) of CON values within 2hr of alcohol exposure, and these reductions persisted through at least 6hr (61 ± 2%, *p*<0.001) and were normalized by 12hr post-exposure (**Figure 2A**). This reduction was mediated, in part, at the transcriptional level because the abundance of transcripts encoding S6K was 89.7% of CON (*p*_adj_=4.28E-12) as determined by whole exome sequencing. With respect to its activated form, phospho-(p)S6K-Threonine(T)389, its levels did not decline as a percentage of total S6K until 12hr post-exposure, when they dropped to 57 ± 1% at 12hr (*p*<0.001) and 59 ± 7% at 18hr (*p*=0.004) post-exposure. However, because of the absolute decline in S6K protein, this represented a 32% decline in the abundance of activated pS6K within 2hr of the alcohol exposure and a 47% decline at 12hr.

**Figure 2.**
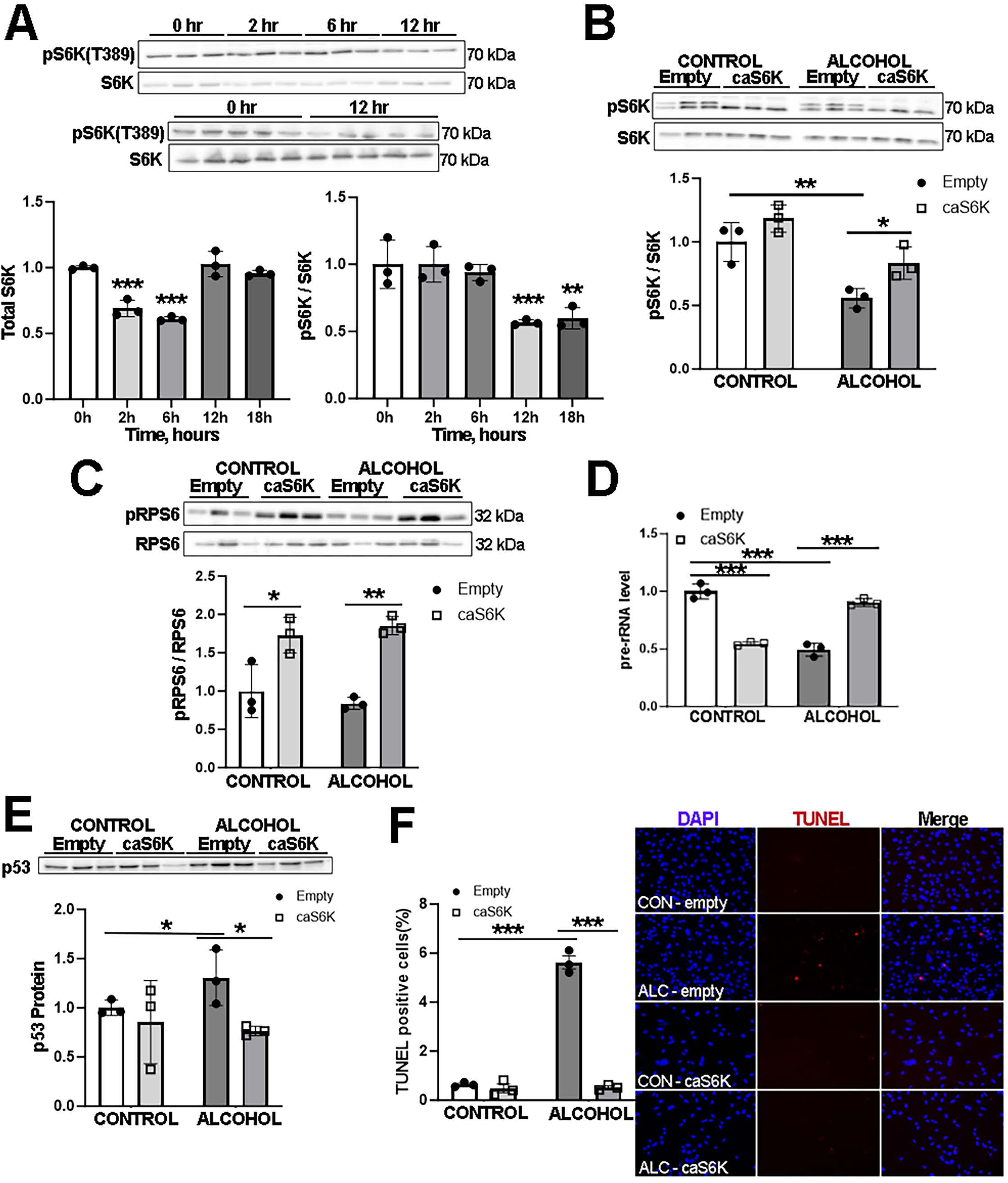
Alcohol reduces the positive effector of ribosome biogenesis S6K. **(A)** Western blot analysis for phospho- and total S6K at the indicated times following alcohol exposure. Total S6K is reduced by 2hr post-exposure; the ratio of pS6K to S6K is reduced by 12hr as compared with control (CON). **(B-F)** Overexpression of gain-of-function caS6K **(B)** enhances its protein abundance in ALC but not CON neural crest, **(C)** elevates the abundance of pRPS6 at 12hr post-exposure, **(D)** normalizes rRNA synthesis in ALC neural crest and reduces it in CON at 2hr post-exposure, **(E)** prevents the elevated p53 content in ALC neural crest at 12hr, and **(F)** prevents alcohol-induced apoptosis, as assessed by TUNEL. All studies use 80mM alcohol. Values are mean ± SD with N=3 independent experiments and analyzed using one-way ANOVA compared with Time 0 (A), or two-way ANOVA compared with CON (B-F). * p<0.05, ** p<0.01, *** p<0.001.

To test the functional involvement of this S6K loss, we transfected neural crest with a constitutively active (ca) version of S6K (caS6K), which contains four gain-of-function mutations that confer rapamycin-resistance [42]. S6K has autocatalytic activity [43, 44], as seen in the elevated levels of pS6K in both CON and ALC transfected cells (**Figure 2B**). A primary target of activated S6K is ribosomal protein S6 (RPS6), and caS6K elevated pRPS6 in both CON (173 ± 19%; *p*=0.015) and ALC cells (185 ± 6%; *p*=0.002; **Figure 2C**); alcohol itself did not affect pRPS6 at this 12hr timepoint (84 ± 9%). The caS6K construct countered the alcohol-mediated reduction in pS6K(T389) (ALC, 55 ± 7%, ALC+caS6K, 83 ± 12%, *p*=0.029; **Figure 2B**) to levels not different from CON (1.00 ± 0.16, *p*>0.10). caS6K also restored *de novo* rRNA synthesis in alcohol-exposed cells (ALC 49 ± 5%, ALC+caS6K 90 ± 3%, *p*<0.001; **Figure 2D**), abrogated alcohol’s activation of p53 (ALC, 155 ± 29%, ALC+caS6K, 106 ± 14%; *p*=0.05; **Figure 2E**), and prevented their apoptosis (**Figure 2F**, ALC, 2.70 ± 0.45%, ALC+caS6K, 0.52± 0.11%; *p* <0.001). In the controls, caS6K unexpectedly reduced rRNA synthesis (CON, 100 ± 6%, CON+caS6K, 54 ± 1%, *p*<0.001), but this had no effect on p53 content or apoptosis. We conclude that alcohol rapidly suppressed S6K and that this contributes to the loss of RBG.

### 3.3. Alcohol activates the upstream TORC1 repressors pTSC2 and pRaptor to reduce TORC1 signaling activity

A primary activator of S6K is the Target of Rapamycin Complex 1 (TORC1). TORC1 is under complex regulation via phosphorylation of its component proteins Raptor and TSC2, which act immediately upstream as AMPK-dependent negative regulators of TORC1 [45,46]. Alcohol increased the content of pRaptor(S792) by 6hr (CON 100 ± 5%, ALC 168 ± 29%; *p*=0.04) and increased pTSC2(S1387) within 2hr of exposure (CON 100 ± 1%, ALC 144 ± 25%; *p*=0.05; **Figure 3A**); pTSC2 was largely normalized by 6hr, whereas pRaptor remained elevated for at least 12hr (ALC 143 ± 2%, *p*<0.001 vs CON). As further evidence that this represented a reduction in TORC1 activity, we found a reduction in a second TORC1 target p4E-BP1(T70) at 12hr post-exposure (CON 111 ± 6%, ALC 85 ± 4%, *p*=0.027; **Figure 3B**), which is otherwise activated by TORC1 to increase protein translation in concert with S6K’s stimulation of RBG [27,28]. We also explored alcohol’s impact on effectors of TORC2 and found no differences with respect to the abundance of pRictor(T1135), Deptor, or pPKCα(S657) at this same 12hr timepoint, suggesting alcohol’s suppression of TORC1 was selective in these neural crest cells (**Figure 3C**).

**Figure 3.**
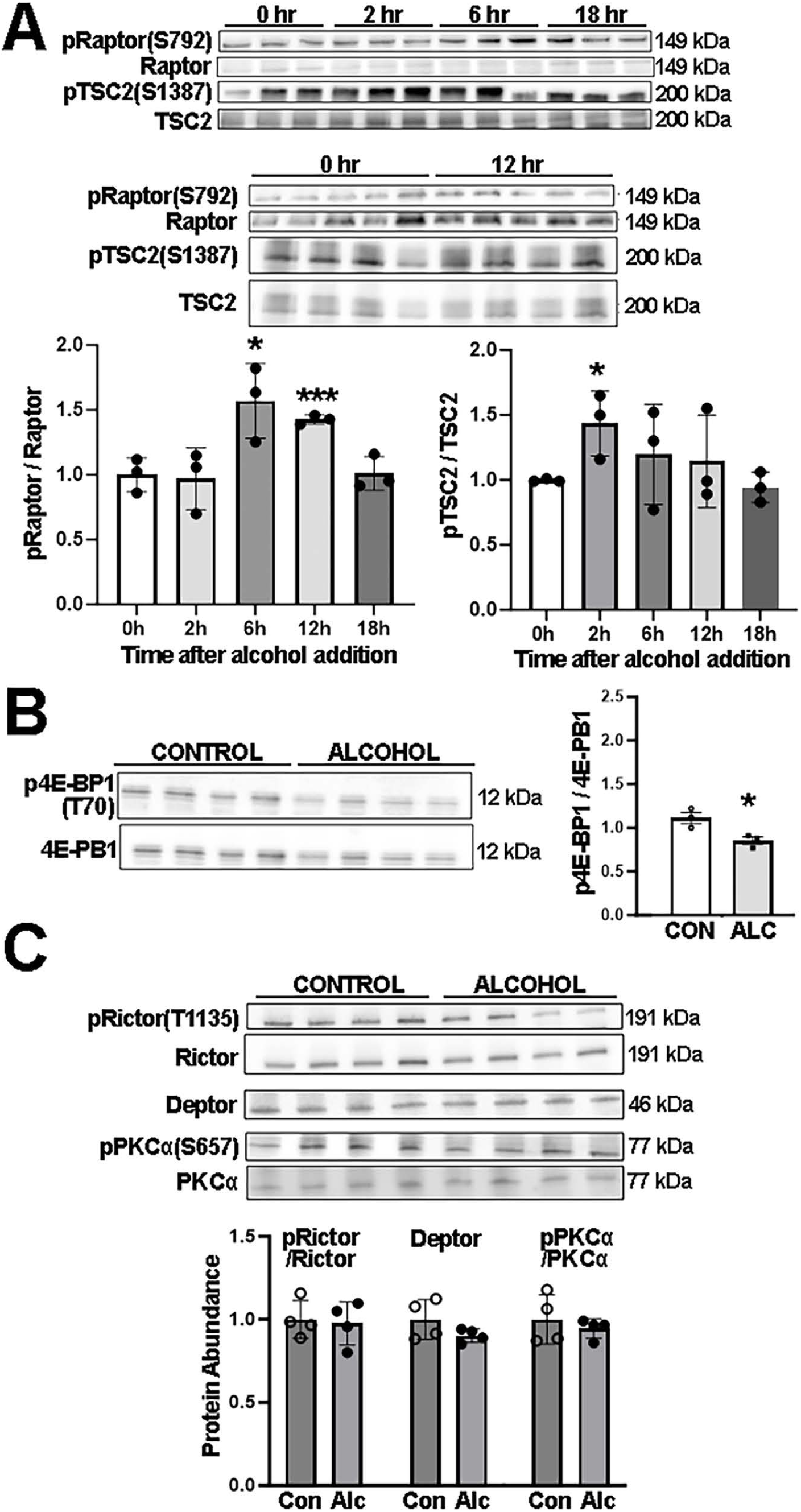
Alcohol represses the upstream S6K effector TORC1 in neural crest. **(A)** Alcohol increases the active phospho-forms of the TORC1 upstream negative effectors pRaptor(S792) and pTSC2(S1387) within 2hr to 6hr of alcohol exposure. **(B)** Alcohol reduces the downstream TORC1 target p4E-BP1 at 12hr post-exposure. **(C)** Alcohol does not affect the abundance of the TORC2 effectors pRictor, Deptor, or PKCα at 12hr post-exposure. Values are mean ± SD with N=3 independent experiments and analyzed using one-way ANOVA (A) or two-tailed t-test (B, C) compared with CON. * p<0.05, ** p<0.01, *** p<0.001.

To explore these findings further, we tested if targeted inhibition of TORC1 signaling, using the highly specific inhibitor rapamycin (Rapa) [47], was sufficient to suppress RBG and induce neural crest apoptosis. In otherwise normal neural crest, rapamycin strongly inhibited the phosphorylation of S6K (CON+Rapa 28 ± 9%, CON 100 ± 15%, p<0.001; **Figure 4A**) and elevated p53 protein (CON+Rapa 159 ± 2%, CON 100 ± 9%, p<0.001; **Figure 4C**), but did not affect rRNA synthesis (**Figure 4B**) or cause apoptosis (**Figure 4D**). In contrast, rapamycin amplified the effects of alcohol exposure to further reduce pS6K (ALC 43 ± 5%, ALC+Rapa 22 ± 3%, p=0.002; **Figure 4A**) and rRNA synthesis (ALC 80 ± 0.3%, ALC+Rapa 59 ± 13%, p=0.035; **Figure 4B**), and further increase p53 protein (ALC 151 ± 21%, ALC+Rapa 175 ± 8%, p=0.050; **Figure 4C**) and apoptosis (ALC 5.19 ±0.37%, ALC+Rapa 7.48± 0.76%, p=0.030; **Figure 4D**).

**Figure 4.**
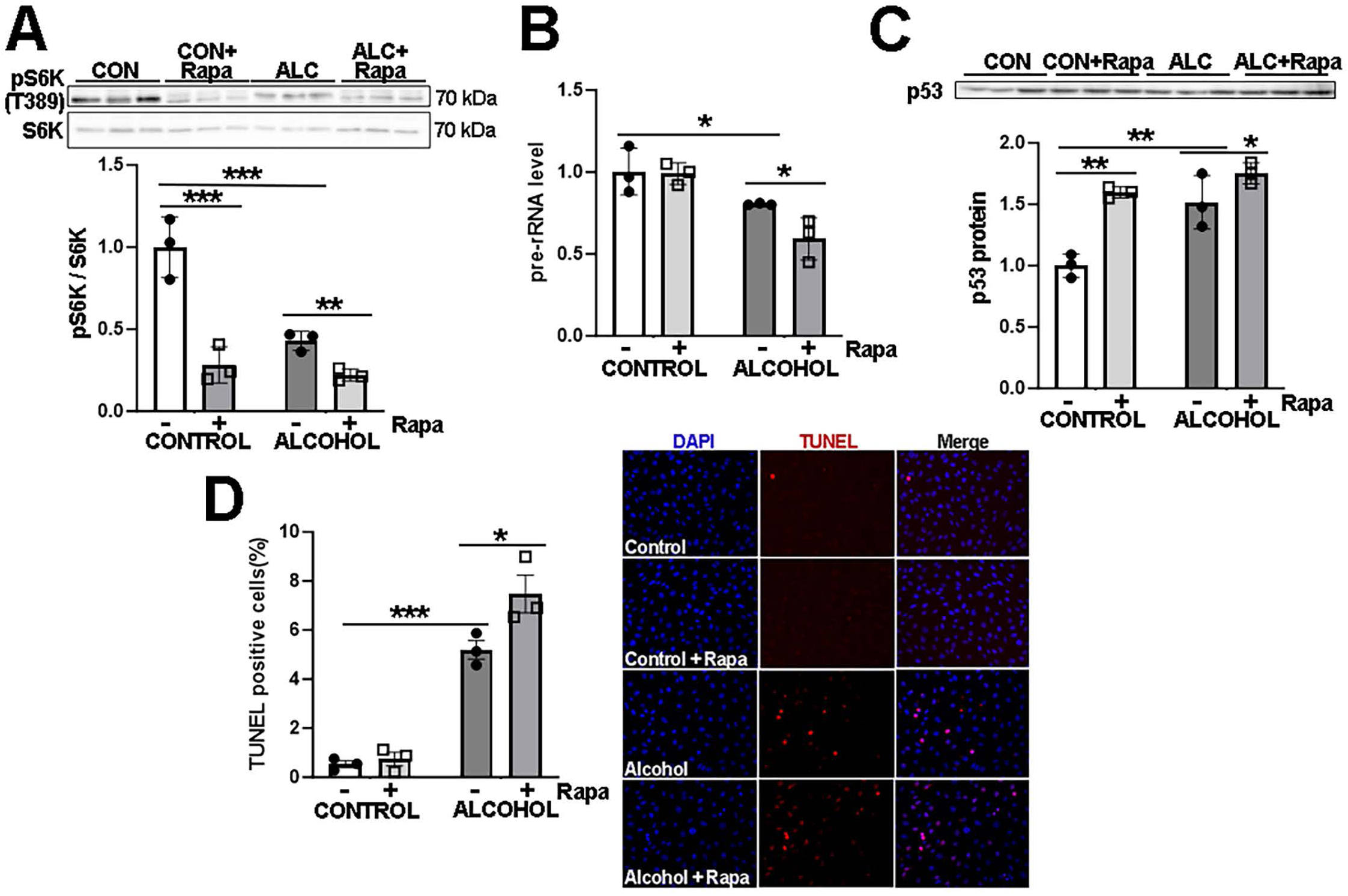
Treatment with the TORC1 inhibitor rapamycin exacerbates alcohol’s impact on rRNA synthesis, p53 stability, and apoptosis in neural crest. **(A)** Rapamycin reduces pS6K in otherwise normal neural crest and further reduces it in ALC neural crest at 12hr post-exposure. **(B)** Rapamycin reduces rRNA synthesis in ALC but not CON neural crest at 2hr post-exposure. **(C)** Rapamycin stabilizes p53 protein in CON and ALC neural crest at 12hr post-exposure. **(D)** Rapamycin causes apoptosis in ALC but not CON neural crest at 16hr post-exposure. Values are mean ± SD with N=3 independent experiments and analyzed using two-way ANOVA compared with CON. * p<0.05, ** p<0.01, *** p<0.001.

We further tested TORC1 contributions functionally by providing L-leucine, which activates TORC1 via the Gag/Raptor pathway [48]. We found that supplemental L-leucine (at 370% of normal cell culture levels) enhanced rRNA synthesis in alcohol-exposed cells (ALC 77 ± 6%, ALC+Leu 143 ± 26%, *p*=0.0042; **Figure 5A**), and abrogated the alcohol-mediated loss of total S6K protein (*p*=0.019) but did not further affect pS6K content (**Figure 5B**). It also normalized p53 levels (*p*<0.001; **Figure 5C**) and prevented their alcohol-induced apoptosis (*p*<0.001; **Figure 5D**). In CON, it increased pS6K levels (*p*=0.021) and rRNA synthesis (CON, 100 ± 5%, CON+Leu, 159 ± 15%, *p*=0.008), and modestly increased p53 content, with no impact on their apoptosis. Taken together, these data suggested that acute alcohol rapidly repressed TORC1 activity and this might be mediated through the activation of Raptor and TSC2. However, it also suggested that suppression of TORC1 was not sufficient, in of itself, to induce the alcohol-mediated apoptosis in neural crest and that additional signals contributed to alcohol’s actions.

**Figure 5.**
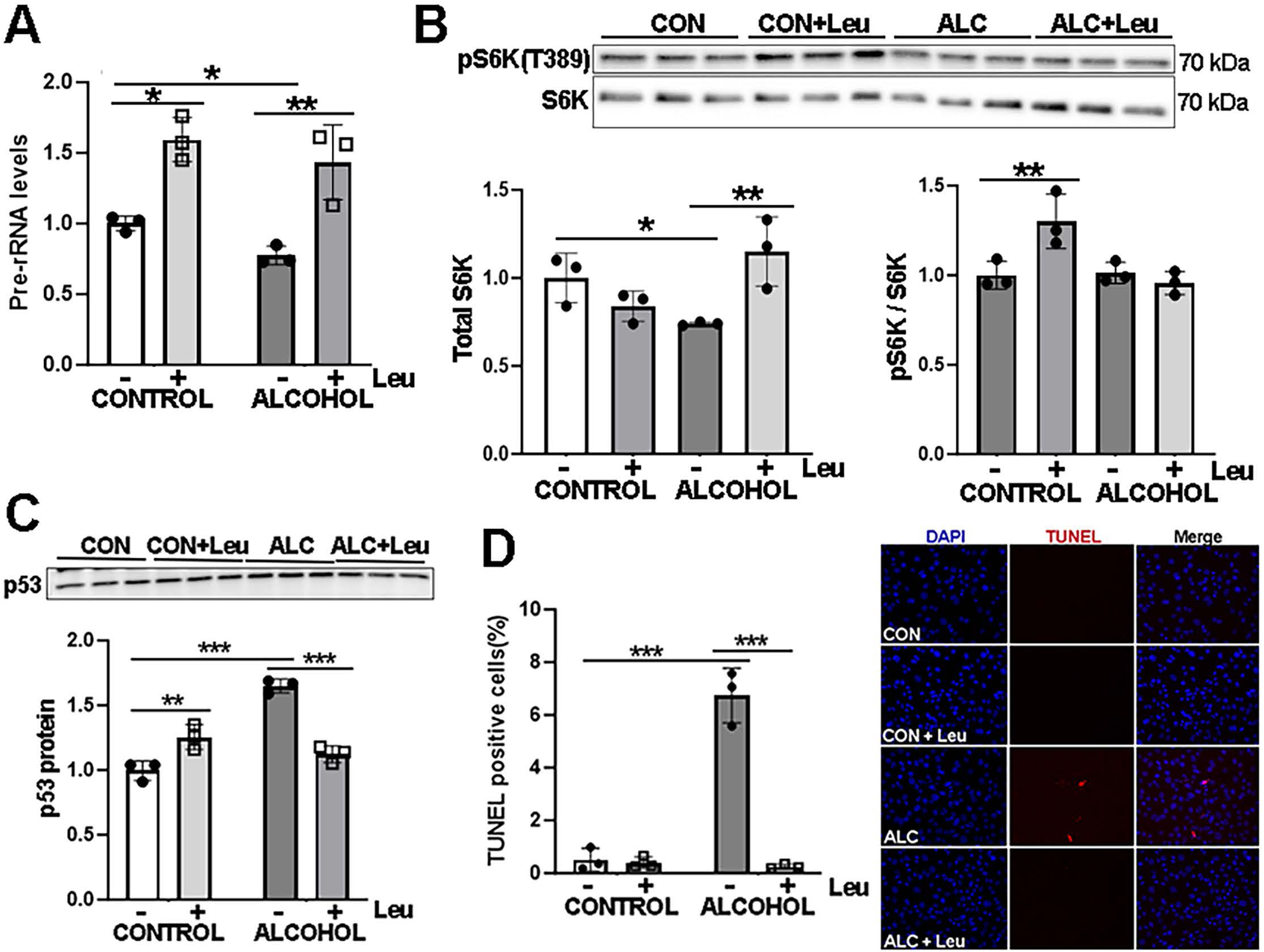
Treatment with the TORC1 activator L-leucine attenuates alcohol’s impact on rRNA synthesis, S6K loss, p53 stability, and apoptosis in neural crest. **(A)** Pretreatment with L-leucine enhances rRNA synthesis in CON and normalizes it in ALC neural crest at 2hr post-exposure. **(B)** L-leucine normalizes total S6K in ALC neural crest at 12hr post-exposure. **(C)** L-leucine reduces p53 protein content in ALC neural crest at 12hr post-exposure. **(D)** L-Leucine normalizes apoptosis in ALC neural crest at 16hr post-exposure. For all, values are mean ± SD with N=3 independent experiments and analyzed using two-way ANOVA compared with CON. * p<0.05, ** p<0.01, *** p<0.001.

### 3.4. Alcohol rapidly activates AMPK signaling in neural crest, and its inhibition prevents alcohol’s suppression of RBG and p53-mediated apoptosis

The canonical repressor of TORC1 activity is the AMP-dependent kinase (AMPK), which inhibits TORC1 via its phosphorylation of TSC2 and Raptor [45,46]. Consistent with its actions upon TSC2, Raptor, and S6K, alcohol exposure caused a rapid elevation in the activated form of AMPK, pAMPK(Thr172), within 2hr of exposure (CON, 100 ± 1%, ALC 143 ± 6%, *p*<0.001; **Figure 6A**). The pAMPK content peaked between 6hr and 12hr (6hr, 188 ± 10%; 12hr, 184 ± 15%; *p*<0.001) and remained elevated through at least 18hr (150 ± 14%, *p*<0.001) following the alcohol exposure. To functionally test AMPK’s role in the loss of S6K and RBG and this p53-mediated apoptosis, we treated cells with the AMPK inhibitor dorsomorphin prior to their alcohol exposure [45,46]. Dorsomorphin attenuated alcohol’s activation of pAMPK(T172) (ALC, 128 ± 7%, ALC+DOR, 89 ± 11%, *p*=0.005) and reversed alcohol’s impact on its downstream targets pRaptor(S792) (ALC, 122 ± 8%, ALC±DOR, 27 ± 3, *p*<0.001) and pS6K(T389) (ALC, 67 ± 6%, ALC+DOR, 135 ± 18%, *p*=0.007; **Figure 6B**). The AMPK inhibitor also normalized *de novo* rRNA synthesis (ALC, 70 ± 4%, ALC±DOR, 97 ± 6%, *p*<0.001; **Figure 6C**), reduced their p53 content (ALC, 128 ± 5%, ALC±DOR, 58 ± 8%; *p*=0.003; **Figure 6D**), and prevented their apoptosis in response to alcohol (ALC 6.74 ± 0.92%, ALC+DOR 0.86 ± 0.06%; *p*<0.001; **Figure 6E**). These data suggested that alcohol caused neural crest apoptosis through the AMPK-mediated inhibition of TORC1 and S6K to suppress their RBG.

**Figure 6.**
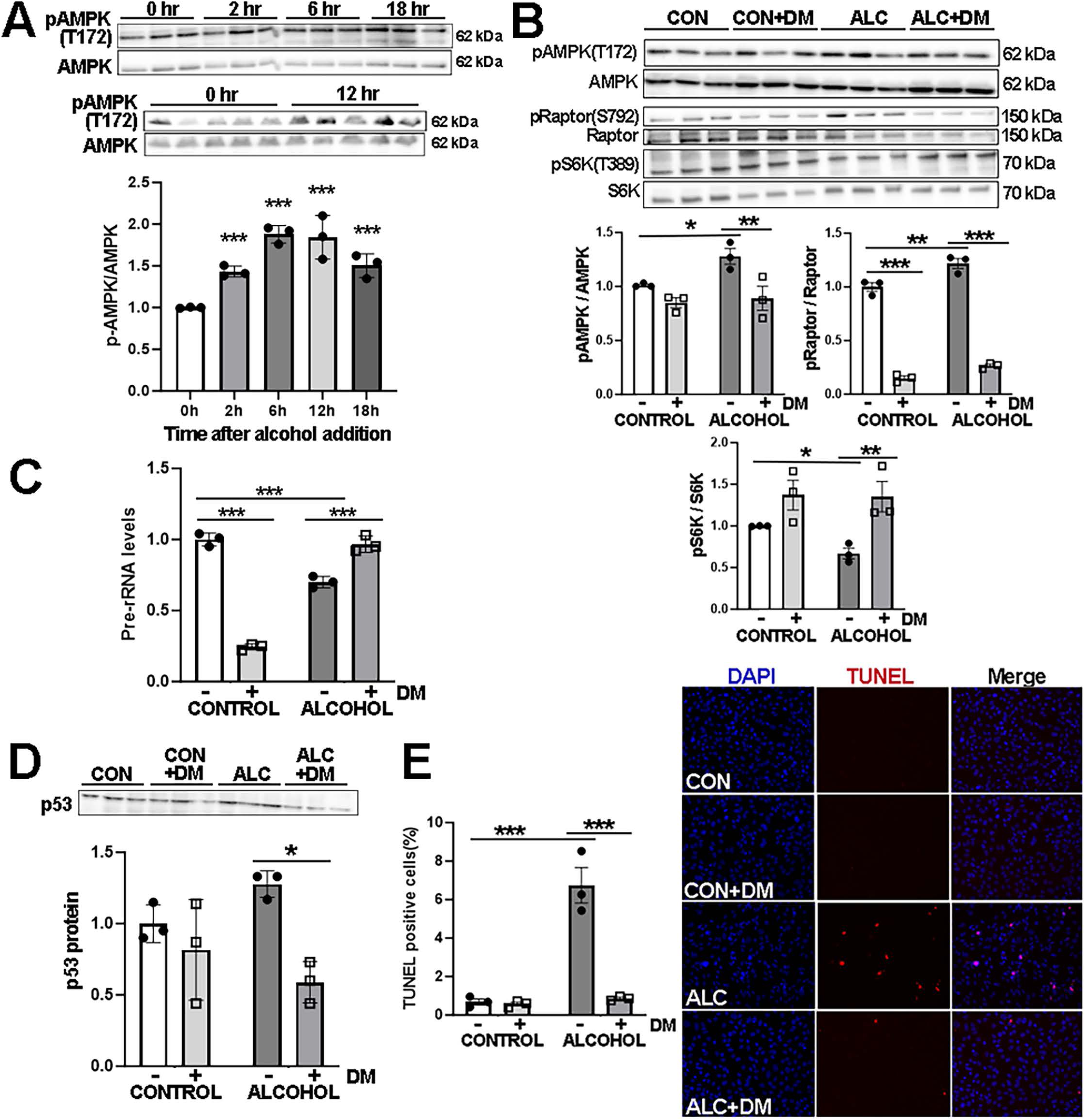
Alcohol rapidly elevates pAMPK in neural crest, and the AMPK inhibitor dorsomorphin counters alcohol’s effects on AMPK-TORC1 signaling, rRNA synthesis, and p53-mediated apoptosis. **(A)** Alcohol exposure increases pAMPK within 2hr of exposure and it remains elevated for at least 18hr. **(B)** The AMPK inhibitor dorsomorphin normalizes pAMPK, pRaptor, and pS6K in ALC neural crest at 12hr post-exposure. **(C-E)** Dorsomorphin also normalizes rRNA synthesis at 2hr **(C)**, p53 protein content at 12hr **(D)**, and apoptosis at 16hr **(E)** in ALC neural crest. Values are mean ± SD with N=3 independent experiments and analyzed using one-way (A) or two-way ANOVA (B-E) compared with CON. * p<0.05, ** p<0.01, *** p<0.001.

However, dorsomorphin is not specific for AMPK and also inhibits the bone morphogenetic protein (BMP) receptors ALK2, ALK3, and ALK6; our transcriptomics reveal that O9-1 cells express ALK2 and ALK3 but not ALK6. We therefore asked if inappropriate activation of pAMPK was sufficient to invoke nucleolar stress and p53-mediated apoptosis in neural crest. Using the AMPK activator A769662 (A76) [45,46,49], in otherwise normal neural crest cells it elevated pAMPK(T172) (CON, 100 ± 2%; CON+A76, 178 ± 3%, *p*=0.010; **Figure 7A**) and reduced p70(T389) (CON, 100 ± 5%; CON+A76, 67 ± 12%; *p*<0.001; **Figure 7A**). AMPK inhibition also strongly reduced rRNA synthesis (CON, 100 ± 5%; CON+A76, 28 ± 2%; *p*<0.001; **Figure 7B**), elevated p53 protein (CON, 100 ± 4%; CON+A76, 174 ± 3%; *p*<0.001; **Figure 7C**), and caused a non-significant rise in apoptosis (CON, 0.54 ± 0.09%; CON+A76, 1.38 ± 0.16%; p=0.053; **Figure 7D**). In alcohol-exposed cells, it further elevated pAMPK(T172) (ALC, 148 ± 17%; ALC+A76, 326 ± 39%, *p*<0.001) and modestly reduced pS6K (ALC, 80 ± 2%; ALC+A76, 65 ± 8%, *p*<0.001; **Figure 7A**), further suppressed rRNA synthesis (ALC, 72 ± 1%; ALC+A76, 50 ± 4%; *p*<0.001; **Figure 7B**), and further elevated p53 protein content (ALC, 128 ± 8%; ALC+A76, 181 ± 7%; *p*<0.001; **Figure 7C**) and apoptosis (ALC, 5.23 ± 47%; ALC+A76, 8.95 ± 0.12%; *p*<0.001; **Figure 7D**).

**Figure 7.**
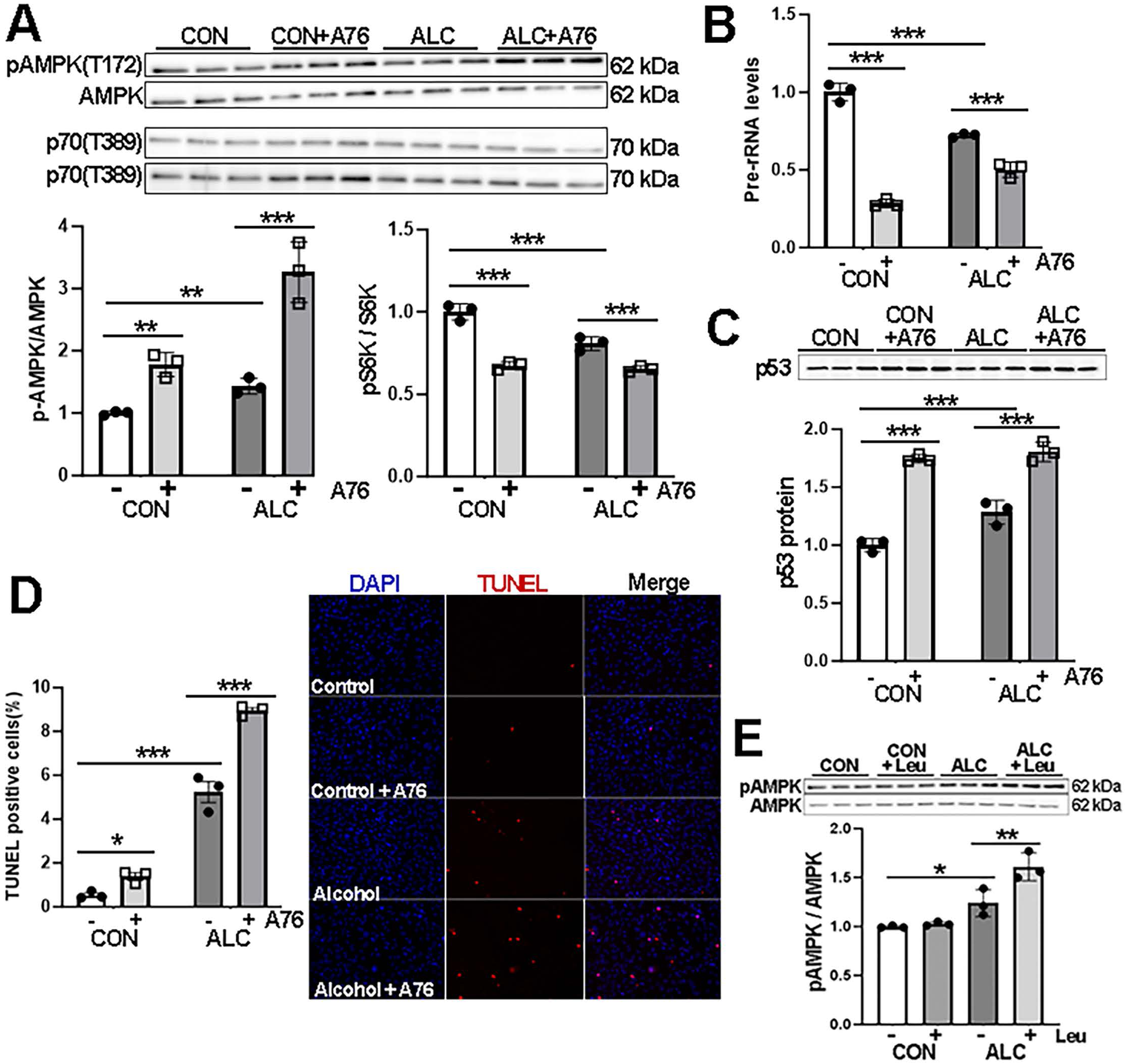
The AMPK activator A769662 further alters AMPK and S6K, suppresses rRNA synthesis and exacerbates p53-mediated apoptosis in both ALC and CON neural crest. **(A)** A769662 further exacerbates pAMPK activation and pS6K in ALC neural crest at 2hr post-exposure, and elevates pAMPK and reduces pS6K in otherwise normal neural crest. **(B-D)** A769662 further impacts rRNA synthesis **(B),** p53 protein content, **(C),** and apoptosis **(D)** in ALC neural crest, and suppresses rRNA synthesis and elevates p53 protein in CON. **(E)** L-Leucine further elevates pAMPK in ALC but not CON cells at 12hr post-exposure. Values are mean ± SD with N=3 independent experiments and analyzed using two-way ANOVA compared with CON. * p<0.05, ** p<0.01, *** p<0.001.

Finally, AMPK and TORC1 operate in opposition to directly repress each other’s activity [45,46], and it was possible that the increased AMPK in response to alcohol represented, not the alcohol-mediated activation of AMPK to suppress TORC1, but rather the alcohol-mediated suppression of TORC1 to elevate AMPK. We explored this again using the TORC1 activator L-leucine. Pretreatment of cells with L-leucine did not affect pAMPK in otherwise normal cells (**Figure 7E**). In the presence of alcohol, L-leucine did not override the activation of pAMPK, and the combination actually elevated pAMPK further (ALC, 1.24 ± 0.14; ALC+Leu, 1.62 ± 0.14, *p*=0.007). This suggested that alcohol’s activation of AMPK was upstream of its suppression of TORC1, and that a reduction in TORC1 was not the primary driver of this increased pAMPK in response to alcohol. Taken together, these data indicate that alcohol’s activation of AMPK is responsible for the S6K-mediated loss of RBG and activation of p53-mediated apoptosis in neural crest.

## 4. Discussion

The most important finding reported here is that alcohol exposure suppressed the TORC1-dependent activity of S6K in cranial neural crest to inhibit ribosome biogenesis, induce nucleolar stress, and effect their p53-mediated apoptosis. Moreover, this suppression of S6K was driven by alcohol’s upstream activation of pAMPK. Counteracting these changes through administration of pS6K gain-of-function or dorsomorphin-mediated inhibition of pAMPK were sufficient to sustain RBG in alcohol’s presence and thus prevent p53 stabilization and apoptosis. Alcohol’s action on this pathway was rapid, with a loss of rRNA synthesis within 1hr and increases in both pAMPK and p53 within 2hr post-exposure, suggesting these represent primary targets of alcohol. We reported previously [24] that pharmacologically relevant alcohol exposures (20-80mM) suppress RBG and induce nucleolar stress in cranial neural crest cells, leading to their p53/MDM2 mediated apoptosis. The rapidity of pAMPK’s activation in response to alcohol, and the ability of interventions targeting AMPK-TORC1-S6K gain- and loss-of-function to alter alcohol’s effects on RBG and p53-mediated apoptosis, suggests that AMPK activation and its downstream suppression of TORC1-S6K are primary effectors of this alcohol-induced apoptosis. These actions of alcohol upon the AMPK-TORC1-S6K pathway to instigate neural crest apoptosis are summarized in **Figure 8**.

**Figure 8.**
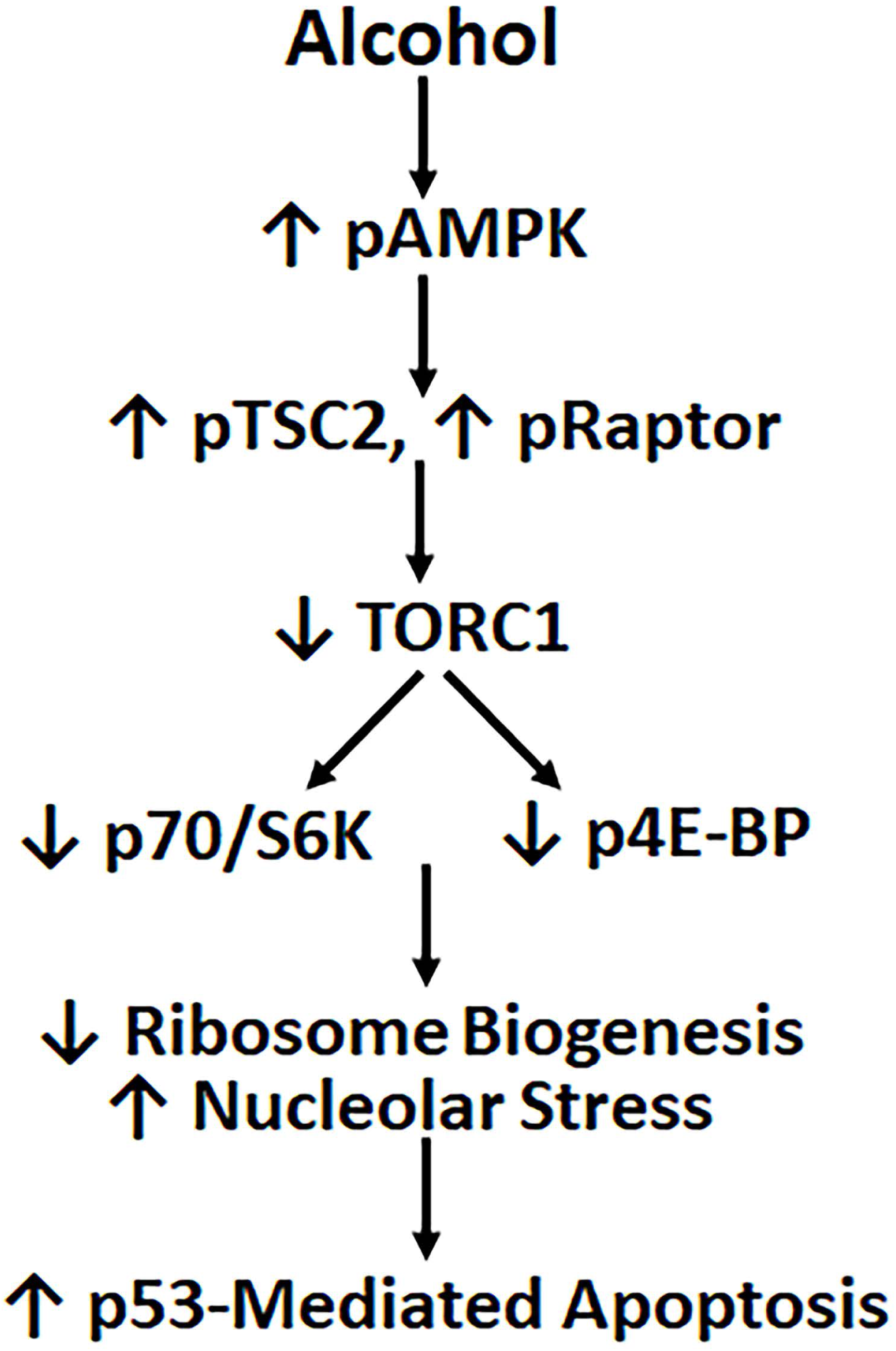
Diagram of alcohol’s proposed mechanism of action in mediating neural crest apoptosis. Data herein and in [24] suggest that alcohol’s activation of pAMPK activates the TORC1 repressors pTSC2 and pRaptor to reduces TORC1’s stimulation of ribosome biogenesis via S6K, resulting in nucleolar stress and the activation of p53/MDM2-mediated apoptosis.

A potential involvement of TORC1 in alcohol-mediated neural crest death has been suggested from prior studies, but those observations have not yet been integrated into the larger, cohesive mechanism investigated here. Specifically, in a zebrafish model of PAE, haploinsufficiency in *Pdgfr* synergizes with alcohol to cause craniofacial deficits, and these are attenuated by treating the embryos with the TORC1 activator L-leucine [50], as TORC1 can operate downstream from PDGFR. Because that study did not directly access TORC1 activity, alcohol’s potential impact on TORC1 in that model is unclear. Acute alcohol exposures reduce TORC1 signaling in adult cerebral cortex [51], similar to its effects here. Although chronic alcohol exposure also reduced p-mTOR in fetal hippocampus [52], this was accompanied by elevated pS6K and p4E-BP1, along with elevated Deptor and Rictor but not Raptor, outcomes distinct from those reported here. It is possible that chronic alcohol exposure led to adaptive or compensatory changes in response to reduced TORC1 or may have dysregulated aspects of that pathway [52]. Our findings are consistent, however, with the increased ULK/ATG signals in alcohol-exposed cells [53,54], as autophagy also increases under TORC1 suppression, and this has been posited to reflect a protective response to alcohol [53,54]; we did not assess autophagy here. Acute alcohol (18-24hr) also suppresses the phosphorylation of the downstream TORC1 targets S6K and 4E-BP1 in SH-SY5Y neuroblastoma cells [55] and C2C12 myocytes [56] and promotes apoptosis in that neuronal model [55]. That TORC1 signaling is indispensable for neural crest is highlighted by studies in which its modulation by genetic [57] or dietary means [58] disrupts craniofacial morphogenesis and alters the facial appearance. Taken together, the critical requirement of neural crest for high rates of RBG to sustain their equally high proliferative rate prior to their migration [25,59,60], and the requirement for TORC1 to sustain S6K activation and RBG [28], could explain how alcohol’s suppression of TORC1 and S6K would enhance their vulnerability to alcohol’s neurotoxicity.

As an anabolic effector of cell growth, TORC1 activity is tightly regulated by both its own direct nutrient sensing and by upstream kinases such as AMPK that further act to integrate metabolic status. As a primary suppressor of TORC1 and RBG [27,45,46], alcohol’s activation of AMPK and its targets TSC2 and Raptor suggest that alcohol reduced RBG via repression of TORC1 signaling. This was further endorsed in that treatment of ALC neural crest with the AMPK antagonist dorsomorphin reversed alcohol’s impact and normalized pAMPK, pS6K, p-mTOR, and pRaptor, and prevented both nucleolar stress and apoptosis. That AMPK inhibition blocked alcohol’s adverse effects indicates that pAMPK participates in their apoptosis. However, this interpretation is tempered in that dorsomorphin also inhibits BMP signaling, and we cannot rule out that it conferred rescue through that interaction. A role for AMPK is further supported in that the AMPK-specific agonist, A769662, mirrored alcohol’s effects on TORC1/S6K/RBG and p53, and exaggerated these outcomes in alcohol-exposed cells. Moreover, neither TORC1 inhibition via rapamycin, nor AMPK activation via A769662 were sufficient to cause apoptosis in otherwise healthy neural crest, and this suggests that alcohol alters signals in addition to pAMPK and TORC1 to induce their nucleolar stress and apoptosis. This is not unexpected as alcohol does not act through a single receptor, but rather binds and modulates the activity of multiple protein targets [61]. For example, we have previously shown that in cranial neural crest alcohol initiates a G-protein-mediated intracellular calcium transient that activates CaMKII and destabilizes nuclear β-catenin to similarly trigger their apoptosis [15,58–60]. Alcohol also has been shown to modulate facial outcome through signals involving sonic hedgehog, retinoids, and oxidative stress, among others [5,6,11–15]. Studies are underway to elucidate these additional alcohol-responsive contributors that further destabilize these neural crest cells and tip them into an apoptotic fate.

Alcohol has been shown to activate AMPK in multiple models including neurons of the hippocampus [65], prefrontal cortex [66], isolated cardiac and myoblast cells [67–69], and hepatocytes [70]. Blockade of AMPK activation prevents neuronal cell death in models of zinc-induced toxicity [71] and Parkinson’s disease [72]. However, how alcohol increases AMPK is currently unclear. AMPK is a master sensor and coordinator of cellular energy status and is activated in response to falling energy status via multiple mechanisms including binding to elevated AMP, carbohydrate binding motifs and, perhaps of relevance to this model, pCAMKK2 in response to intracellular calcium signals [45,46,73]. There is debate whether AMPK activation in response to alcohol represents a beneficial or deleterious effect [74], and studies are contradictory in this regard. AMPK can increase glycolytic flux in response to hypoxia, as well as acting to limit energy-demanding processes such as RBG [45,46]. This latter may be especially detrimental to cranial neural crest given their absolute dependence upon RBG as evidenced in the genetic disorders known as ribosomopathies (i.e. Treacher-Collins, Diamond-Blackfan Anemia), in which RBG loss-of-function causes p53-mediated neural crest losses and facial deficits that parallel those of PAE including flattened philtrum, thin upper lip, epicanthal folds, flattened midface, and micrognathia [1–6,34–36]. What is unclear is why alcohol’s suppression of RBG invokes a nucleolar stress instead of a ‘soft landing’ or adaptive response that protects against p53 activation and MDM2/p53-mediated apoptosis, and investigations into this are underway.

In conclusion, pharmacologically relevant alcohol exposures significantly enhance the phosphorylation of AMPK in cranial neural crest and inhibit the downstream TORC1 signaling pathway to reduce activated S6K and 4E-BP1. This is accompanied by a suppression of *de novo* rRNA synthesis, induction of nucleolar stress, and stabilization of p53 to initiate their apoptosis. These findings underscore the pivotal role of the AMPK/TORC1 signaling axis as a novel mechanism in mediating alcohol’s deleterious effects on neural crest development and survival.

## Acknowledgements

We are grateful to Josh Bausch for early work on this project, and to Thomas Wilkie for helpful discussions and assistance.

## Author Contributions

YH - Conceptualization, methodology, formal analysis, investigation, writing original draft, writing edit and review, visualization; GRF - Conceptualization, methodology, investigation, writing edit and review; SMS - Conceptualization, writing original draft, writing edit and review, visualization, administration, funding.

## References

1. Hoyme HE, Kalberg WO, Elliott AJ, Blankenship J, Buckley D, Marias AS, et al. 2016. Updated clinical guidelines for diagnosing fetal alcohol spectrum disorders. Pediatrics 138:e201542568.

2. Suttie M, Foroud T, Wetherill L, Jacobson JL, Molteno CD, Meintjes EM, Hoyme HE, Khaole N, Robinson LK, Riley EP, Jacobson SW, Hammond P. 2013. Facial dysmorphism across the fetal alcohol spectrum. Pediatrics. 131:e779–88.

3. Foroud T, Wetherill L, Vinci-Booher S, Moore ES, Ward RE, Hoyme HE, Robinson LK, Rogers J, Meintjes EM, Molteno CD, Jacobson JL, Jacobson SW. 2012. Relation over time between facial measurements and cognitive outcomes in fetal alcohol-exposed children. Alcohol Clin Exp Res 36:1634–1646.

4. Lipinski RJ, Hammond P, O’Leary-Moore SK, Ament JJ, Pecevich SJ, Jiang Y, Budin F, Parnell SE, Suttie M, Godin EA, Everson JL, Dehart DB, Oguz I, Holloway HT, Styner MA, Johnson GA, Sulik KK. 2012. Ethanol-induced face-brain dysmorphology patterns are correlative and exposure-stage dependent. PLoS One. 7(8):e43067.

5. Smith SM, Garic A, Flentke GR, Berres ME. 2014. Neural crest development in fetal alcohol syndrome. Birth Defects Res C Embryo Today. 102:210–20.

6. Chen SY, Kannan M. 2023. Neural crest cells and fetal alcohol spectrum disorders: Mechanisms and potential targets for prevention. Pharmacol Res. 194:106855.

7. Bhatt S, Diaz R, Trainor PA. 2013. Signals and switches in mammalian neural crest cell differentiation. Cold Spring Harb Perspect Biol. 5(2):a008326.

8. Cartwright MM, Tessmer LA, Smith SM. 1998. Ethanol-induced neural crest apoptosis is coincident with their endogenous death but is mechanistically distinct. Alcohol Clin Exp Res 22:142–149.

9. Dunty WC Jr, Chen SY, Zucker RM, Dehart DB, Sulik KK. 2001. Selective vulnerability of embryonic cell populations to ethanol-induced apoptosis: implications for alcohol-related birth defects and neurodevelopmental disorder. Alcohol Clin Exp Res. 36:1340–1354.

10. Fan H, Li Y, Yuan F, Lu L, Liu J, Feng W, Zhang HG, Chen SY. 2022. Up-regulation of microRNA-34a mediates ethanol-induced impairment of neural crest cell migration in vitro and in zebrafish embryos through modulating epithelial-mesenchymal transition by targeting Snail1. Toxicol Lett. 358:17–26.

11. Mazumdar R, Eberhart JK. 2023 Loss of nicotinamide nucleotide transhydrogenase sensitizes embryos to ethanol-induced neural crest and neural apoptosis via generation of reactive oxygen species. Front Neurosci. 17:1154621.

12. Burton DF, Boa-Amponsem OM, Dixon MS, Hopkins MJ, Herbin TA, Toney S, Tarpley M, Rodriguez BV, Fish EW, Parnell SE, Cole GJ, Williams KP. 2022. Pharmacological activation of the Sonic hedgehog pathway with a Smoothened small molecule agonist ameliorates the severity of alcohol-induced morphological and behavioral birth defects in a zebrafish model of fetal alcohol spectrum disorder. J Neurosci Res.100(8):1585–1601.

13. Petrelli B, Bendelac L, Hicks GG, Fainsod A. 2019. Insights into retinoic acid deficiency and the induction of craniofacial malformations and microcephaly in fetal alcohol spectrum disorder. Genesis. 57:e23278.

14. Chen X, Liu J, Chen SY. 2013. Over-expression of Nrf2 diminishes ethanol-induced oxidative stress and apoptosis in neural crest cells by inducing an antioxidant response. Repro Tox. 42:102–109.

15. Flentke GR, Garic A, Amberger E, Hernandez M, Smith SM. 2011. The calcium-mediated repression of β-catenin and its transcriptional signaling mediates neural crest cell death in an avian model of fetal Alcohol Syndrome. Birth Defects Res A. 91:591–602.

16. Fish EW, Tucker SK, Peterson RL, Eberhart JK, Parnell SE. 2021. Loss of tumor protein 53 protects against alcohol-induced facial malformations in mice and zebrafish. Alcohol Clin Exp Res. 45(10):1965–1979.

17. Flentke GR, Baulch JW, Berres ME, Garic A, Smith SM. 2019. Alcohol-mediated calcium signals dysregulate pro-survival Snai2/PUMA/Bcl2 networks to promote p53-mediated apoptosis in avian neural crest progenitors. Birth Defects Res. 111:686–699.

18. Berres ME, Garic A, Flentke GR, Smith SM. 2017. Transcriptome profiling identifies ribosome biogenesis as a target of alcohol teratogenicity and vulnerability during early embryogenesis. PLoS ONE e0169351.

19. Li F, Lin J, Li T, Jian J, Zhang Q, Zhang Y, Liu X, Li Q. 2022. Rrn3 gene knockout affects ethanol-induced locomotion in adult heterozygous zebrafish. Psychopharmacology (Berl) 239(2):621–630.

20. Green ML, Singh AV, Zhang Y, Nemeth KA, Sulik KK, Knudsen TB. 2007. Reprogramming of genetic networks during initiation of the Fetal Alcohol Syndrome. Dev Dyn. 236:613–631.

21. Downing C, Balderrama-Durbin C, Kimball A, Biers J, Wright H, Gilliam D, Johnson TE. 2012. Quantitative trait locus mapping for ethanol teratogenesis in BXD recombinant inbred mice. Alcohol Clin Exp Res. 36:1340–1354.

22. Garic A, Berres ME, Smith SM. 2014. High-throughput transcriptome sequencing identifies candidate genetic modifiers of vulnerability to fetal alcohol spectrum disorders. Alcohol Clin Exp Res. 38:1874–1882.

23. Huang Y, Flentke GR, Rivera OC, Saini N, Mooney SM, Smith SM. 2024. Alcohol exposure induces nucleolar stress and apoptosis in mouse neural stem cells and late-term fetal brain. Cells 13(5):440.

24. Flentke GR, Wilkie TE, Baulch J, Huang Y, Smith SM. 2024. Alcohol exposure suppresses ribosome biogenesis and causes nucleolar stress in cranial neural crest cells.. PLoS One. 19(6):e0304557.

25. Warner JR, Vilardell J, Sohn JH. 2001. Economics of ribosome biosynthesis. Cold Spr Harbor Symp Quant Biol LXVI:567–574.

26. Nomura M. 2001. Ribosomal RNA genes, RNA polymerases, nucleolar structures, and synthesis of rRNA in the yeast Saccharomyces cerevisiae. Cold Spring Harb Symp Quant Biol 66:555–565.

27. Dibble CC, Manning BD. Signal integration by mTORC1 coordinates nutrient input with biosynthetic output. Nature Cell Biol. 15:555–564.

28. Chauvin C, Koka V, Nouschi A, Mieulet V, Hoareau-Aveilla C, Dreazen A, Cagnard N, Carpentier W, Kiss T, Meyuhas O, Pende M. 2014. Ribosomal protein S6 kinase activity controls the ribosome biogenesis transcriptional program. Oncogene. 33:474–83.

29. Boulon S, Westman BJ, Hutten S, Boisvert F-M, Lamond AI. 2010. The nucleolus under stress. Mol Cell 40:216–227.

30. Lafita-Navarro MC, Conacci-Sorrell M. 2023. Nucleolar stress: From development to cancer. Semin Cell Dev Biol. 136:64–74.

31. Chakraborty A, Uechi T, Kenmochi N. 2011. Guarding the ‘translation apparatus’: defective ribosome biogenesis and the p53 signaling pathway. WIREs RNA 2:507–522.

32. Deisenroth C, Franklin DA, Zhang Y. 2016. The evolution of the ribosomal protein-MDM2-p53 pathway. Cold Spring Harb Perspect Med. 6(12):a026138.

33. Daftuar L, Zhu Y, Jacq X, Prives C. 2013. Ribosomal proteins RPL37, RPS15, and RPS20 regulate the Mdm2-p53-MdmX network. PLoS ONE 8:e68667.

34. Yelick PC, Trainor PA. 2015. Ribosomopathies: global process, tissue specific defects. Rare Dis 3(1):e1025185

35. Trainor PA. 2014. Craniofacial birth defects: the role of neural crest cells in the etiology and pathogenesis of Treacher Collins syndrome and the potential for prevention. Am J Med Genet Part A 152A:2984–2994.

36. Kang J, Brajanovski N, Chan KT, Xuan J, Pearson RB, Sanij E. 2021. Ribosomal proteins and human diseases: molecular mechanisms and targeted therapy. Signal Transduct Target Ther. 6:323.

37. Ishii M, Arias AC, Liu L, Chen YP, Bronner ME, Maxson RE. 2012. A stable cranial neural crest cell line from mouse. Stem Cells Dev. 21:3069–80.

38. Nguyen BH, Ishii M, Maxson RE, Wang J. 2018. Culturing and manipulation of O9-1 neural crest cells. J Vis Exp. 140:58346.

39. Wang J, Xiao Y, Hsu CW, Martinez-Traverso IM, Zhang M, Bai Y, Ishii M, Maxson RE, Olson EN, Dickinson ME, Wythe JD, Martin JF. 2016. Yap and Taz play a crucial role in neural crest-derived craniofacial development. Development. 143:504–515.

40. Pascual F, Icyuz M, Karmaus P, Brooks A, Van Gorder E, Fessler MB, Shaw ND. 2023. Cholesterol biosynthesis modulates differentiation in murine cranial neural crest cells. Sci Rep. 13:7073.

41. Bustin SA, Benes V, Garson JA, Hellemans J, Huggett J, Kubista M, Mueller R, Nolan T, Pfaffl MW, Shipley JL, Vandesompele J, Wittwer CT. 2009. The MIQE guidelines - minimum information for publication of quantitative real-time PCR experiments. Clin Chem 55:611–622.

42. Lee-Fruman KK, Kuo CJ, Lippincott J, Terada N, Blenis J. 1999. Characterization of S6K2, a novel kinase homologous to S6K1. Oncogene. 18:5108–5114.

43. McCoy CE, Campbell DG, Deak M, Bloomberg GB, Arthur JSC. 2005. MSK1 activity is controlled by multiple phosphorylation sites. Biochem J. 387(Pt 2):507–517; autocatalytic S6K

44. Deak M, Clifton AD, Lucocq LM, Alessi DR. 1998. Mitogen- and stress-activated protein kinase-1 (MSK1) is directly activated by MAPK and SAPK2/p38, and may mediate activation of CREB. EMBO J. 17:4426–4441. autocatalytic S6K

45. Steinberg GR, Hardie DG. 2023. New insights into activation and function of the AMPK. Nat Rev Mol Cell Biol. 24:255–272.

46. Gonzalez A, Hall MN, Lin S-C, Hardie DG. 2020. AMPK and TOR: The Yin and Yang of Cellular Nutrient Sensing and Growth Control. Cell Metabolism 31:472–492.

47. Romanyuk N, Sintakova K, Arzhanov I, Horak M, Gandhi C, Jhanwar-Uniyal M, Jendelova P. 2024. mTOR pathway inhibition alters proliferation as well as differentiation of neural stem cells. Front Cell Neurosci 18:1298182.

48. Dodd KM, Tee AR. 2012. Leucine and mTORC1: a complex relationship. Am J Physiol Endocrin Metab. 302:E1329–1342.

49. Sanders MJ, Ali ZS, Hegarty BD, Heath R, Snowden MA, Carling D. 2007. Defining the mechanism of activation of AMP-activated protein kinase by the small molecule A-769662, a member of the thienopyridone family. J Biol Chem. 282:32539

50. McCarthy N, Wetherill L, Lovely CB, Swartz ME, Foroud TM, Eberhart JK. 2013. Pdgfra protects against ethanol-induced craniofacial defects in a zebrafish model of FASD. Development. 140:3254–3265.

51. Li Q, Ren J. 2007. Chronic alcohol consumption alters mammalian target of rapamycin (mTOR), reduces ribosomal p70S6 kinase and p4E-BP1 levels in mouse cerebral cortex. Exp Neurol 204:840–844.

52. Lee J, Lunde-Young R, Naik V, Ramirez J, Orzabal M, Ramadoss J. 2020. Chronic binge alcohol exposure during pregnancy alters mTOR system in rat fetal hippocampus. Alcohol Clin Exp Res. 44:1329–1336.

53. Luo J. Autophagy and ethanol neurotoxicity. 2014. Autophagy. 10:2099–2108.

54. Chen G, Ke Z, Xu M, Liao M, Wang X, Qi Y, Zhang T, Frank JA, Bower KA, Shi X, Luo J. 2012. Autophagy is a protective response to ethanol neurotoxicity. Autophagy. 8:1577–1589.

55. Sangaunchom P, Dharmasaroja P. 2020. Caffeine potentiates ethanol-induced neurotoxicity through mTOR/p70S6K/4E-BP1 inhibition in SH-SY5Y cells. Int J Toxicol 39:131–140.

56. Hong-Brown LQ, Brown CR, Kazi AA, Navaratnarajah M, Lang CH. 2012. Rag GTPases and AMPK/TSC2/Rheb mediate the differential regulation of mTORC1 signaling in response to alcohol and leucine. Am J Physiol Cell Physiol 302:C1557–C1565.

57. Nie X, Zheng J, Ricupero CL, He L, Jiao K, Mao JJ. 2018. mTOR acts as a pivotal signaling hub for neural crest cells during craniofacial development. PLoS Genet 14:e1007491.

58. Xie M, Kaiser M, Gershtein Y, Schnyder D, Deviatiiarov R, Gazizova G, Shagimardanova E, Zikmund T, Kerckhofs G, Ivashkin E. 2024. The level of protein in the maternal murine diet modulates the facial appearance of the offspring via mTORC1 signaling. Nat Comm 15:1–15.

59. Burstyn-Cohen T, Kalcheim C. 2002. Association between the cell cycle and neural crest delamination through specific regulation of G1/S transition. Dev Cell 3:383–395.

60. Ridenour DA, McLennan R, Teddy JM, Semerad CL, Haug JS, Kulesa PM. 2014. The neural crest cell cycle is related to phases of migration in the head. Development. 141:1095–103.

61. Dwyer DS, Bradley RJ. 2000. Chemical properties of alcohols and their protein binding sites. Cell Mol Leif Sci. 57:265–275.

62. Garic-Stankovic A, Hernandez M, Flentke GR, Smith SM. 2006. Structural constraints for alcohol-stimulated Ca2+ release in neural crest, and dual agonist/antagonist properties of n-octanol. Alcohol Clin Exp Res 30:552–559.

63. Garic-Stankovic A., Hernandez MA, Flentke GR, Debelak-Kragtorp KA, Armant DR, Smith SM. 2005. Ethanol triggers neural crest apoptosis thru the selective activation of a pertussis toxin-sensitive G-protein and a phospholipase Cβ-dependent Ca2+ transient. Alcohol Clin Exp Res 29:1237–1246.

64. Flentke GR, Klingler RH, Tanguay RL, Carvan MJ 3rd, Smith SM. 2014. An evolutionarily-conserved mechanism of calcium-dependent neurotoxicity in a zebrafish model of FASD. Alcohol Clin Exp Res. 38:1255–1265.

65. Srivastava VK, Hiney JK, Dees WL. 2018. Alcohol delays the onset of puberty in the female rat by altering key hypothalamic events. Alcohol Clin Exp Res 42:1166–1176.

66. Xu S, Jeong SJ, Li G, Koo JW, Kang UG. 2020. Repeated ethanol exposure influences key enzymes in cholesterol and lipid homeostasis via the AMPK pathway in the rat prefrontal cortex. Alcohol 85:49–56.

67. Hong-Brown LQ, Brown CR, Kazi AA, Huber DS, Pruznak AM, Lang CH. 2010. Alcohol and PRAS40 knockdown decrease mTOR activity and protein synthesis via AMPK signaling and changes in mTORC1 interaction. J Cell Biochem 109:1172–1184.

68. Hong-Brown LQ, Brown CR, Kazi AA, Navaratnarajah M, Lang CH. 2012. Rag GTPases and AMPK/TSC2/Rheb mediate the differential regulation of mTORC1 signaling in response to alcohol and leucine. Am J Physiol Cell Physiol 302:C1557–C1565.

69. Kandadi MR, Hu N, Ren J. 2013. ULK1 plays a critical role in AMPK-mediated myocardial autophagy and contractile dysfunction following acute alcohol challenge. Curr Pharm Des 19:4874–4887.

70. Yuan F, Xu Y, You K, Zhang J, Yang F, Li YX. 2021. Calcitriol alleviates ethanol-induced hepatotoxicity via AMPK/mTOR-mediated autophagy. Arch Biochem Biophys 697:108694.

71. Eom JW, Lee JM, Koh JY, Kim YH. 2016. AMP-activated protein kinase contributes to zinc-induced neuronal death via activation by LKB1 and induction of Bim in mouse cortical cultures. Mol Brain 9:14.

72. Xu Y, Liu C, Chen S, Ye Y, Guo M, Ren Q, Liu L, Zhang H, Xu C, Zhou Q, Huang S, Chen L. 2014. Activation of AMPK and inactivation of Akt result in suppression of mTOR-mediated S6K1 and 4E-BP1 pathways leading to neuronal cell death in in vitro models of Parkinson’s disease. Cell Signal 26:1680–1689.

73. Marcelo KL, Means AR, York B. 2016. The Ca2+/calmodulin/CaMKK2 axis: nature’s metabolic CaMshaft. Trends Endocrin Metab 27:706–718.

74. Naseer M, Ullah I, Narasimhan M, Lee H, Bressan R, Yoon G, Yun D, Kim M. 2014. Neuroprotective effect of osmotin against ethanol-induced apoptotic neurodegeneration in the developing rat brain. Cell Death Dis 5:e1150–e1150.

